# Virus infections in *Varroa destructor*-resistant honeybees

**DOI:** 10.1101/2020.10.16.342246

**Authors:** Melissa A. Y. Oddie, Sandra Lanz, Bjørn Dahle, Orlando Yañez, Peter Neumann

## Abstract

Populations of European honeybee subspecies, *Apis mellifera*, have the ability to adapt naturally to the ectoparasitic mite, *Varroa destructor*. It is possible that a tolerance to mite-vectored viruses may contribute to colony survival. If this is the case, surviving populations should show lower virus titers and prevalence compared to susceptible populations. Here, we investigated the prevalence and titers of 10 viruses, some known to be associated with *V. destructor*, in adult workers and pupae as well as mites. Samples were collected from both a mite-surviving and mite-susceptible honeybee population in Norway. Surviving colonies had a lower prevalence of a key virus (DWV-A) associated with *V. destructor* in individual adult bees sampled, and generally lower titers of this virus in mite infested pupae and mites within the colonies when compared to sympatric, susceptible controls. However, these surviving colonies also displayed higher prevalence and titers of two viruses not associated with *V. destructor* (BQCV & LSV1). The results of this study therefore suggest that general tolerance to virus infections is unlikely to be a key mechanism for natural colony survival in Norway, but evidence may point to mite control as a predominant mechanism.

## Introduction

Certain honeybee pathogens, such as DWV and its recombinants, which in the past were largely benign [1], have developed novel transmission pathways through the invasive ectoparasitic mite Varroa destructor [2] that bolster titers to the point of reducing the longevity of individual bees. This leads to colony weakening and collapse in many cases. [3,4]. As a result, *Varroa destructor*, which made a host shift form the Eastern honeybee *Apis cerana* to the Western honeybee *Apis mellifera*, is one of the most serious threats to Western honeybees, both domestic and wild [3–5]. Due to the limited defense mechanisms of Western honeybees, their unhindered population growth can cause numbers to reach devastating levels in late summer and autumn, when the bees required for winter hibernation are being reared [3,6]. It is known now that Western honeybee populations left untreated for at least five years have the potential to develop the ability to manage *V. destructor* infestations without the need for human-mediated mite control [7,8]. Previous evidence has pointed to several mechanisms: both mechanical, such as reduced post-capping period [9] and potential changes in brood volatiles [10], and behavioral, such as grooming [11], brood removal [12,13] and brood cell recapping [14]. One of the most prominent traits that has been detected in surviving bees, regardless of the mechanisms identified, is suppressed mite reproduction (SMR) [7,15], signified by lower reproductive output on average per foundress each reproductive cycle. SMR has been recorded not only in Africanized bee populations surviving *V. destructor* [16], but in the “more susceptible” European populations as well, including the Primorsky bees originating in Russia [17], the Gotland bees in Sweden, the Avignon bees in France [15,18] and a population in the Oslo region of Norway [8]. It has been proposed that viral tolerance is also a contributing mechanism, possibly due to changes in the dominant viral strain [19,20] or in the bees or mites themselves [21].

A domestic population of surviving bees in Norway [8] has been used commercially since before the introduction of *V. destructor* into the local area approximately 30 years ago. Selection efforts employed by the beekeeper managing the population included the monitoring of high honey productivity, and this trait was preserved along with the development of the mite-surviving adaptations that stemmed from the lack of treatment and subsequent selective pressure [8]. Evidence has been gathered that suggest these bees possess mechanisms that focus on controlling the parasite directly, through SMR [7,8,14]. However, tolerance to viruses is an additional possibility when considering reasons for survivability [21]. In addition, tracking bee viral loads for viruses unlinked to V. destructor may provide us with insight into the potential weaknesses of a rapidly adapted bee population. This study investigated potential viral tolerances and susceptibilities in a population of bees left regularly untreated for V. destructor (the surviving population) by comparing their viral profiles with regularly treated, *V. destructor*-susceptible controls in the same area. Viruses known to be associated with *V. destructor* infestations (DWV, SBPV) [22] as well as eight other viruses (BQCV, KBV, LSV1, LSV2, SBV, CBPV, IAPV, ABPV) were monitored in worker bees throughout the active season. DWV was monitored in both developing pupae and mites contained in their brood cells. The goal was to examine viral titer levels and gather information on the bees’ potential ability to tolerate viral spread by V. destructor, or else reinforce previous findings that SMR and mite population control (and the subsequent decline in high-titer prevalence of DWV) is the central mechanism employed by Norwegian bees to achieve natural survivability. If viral prevalence and titer in surviving populations are high, we can assume viral tolerance plays a role. If only titer appears lower, we can assume an internal mechanism to fight infection and if prevalence is lower, we can infer that there is a mechanism that limits the spread of the virus, such as reducing mite loads. The other aim of the study was to measure viral prevalence and titer in viruses unassociated with *V. destructor* to detect potential susceptibility in the surviving population to viruses that would, in un-adapted populations, not pose much of a problem. If titers or prevalence of any of these viruses are higher than in sympatric controls we could conclude that heightened susceptibilities may be present, possibly due to a genetic bottlenecking during the natural selection process.

## Materials and Methods

### Sample collection and Varroa destructor infestation rates

In autumn 2013, spring and summer 2014, samples were taken in Østlandet, south-eastern Norway from local queenright *A. mellifera* colonies surviving *V. destructor* infestation without treatment for at least 16 years (*N*=32, 3 apiaries), as well as colonies, regularly treated until 2011/2012 with oxalic acid and/or drone brood removal, (*N*=69, 7 apiaries). Adult workers were collected from outer frames inside the hive. Phoretic mites were sampled using routine washing methods (~100-400 bees, [23]). Infested pupae were sampled, and their mites collected and stored. All samples were transported on ice to Bern, Switzerland for preserving [24] and then stored at −80°C until processing.

### Sample selection and analytic approach

#### Pooled samples

100 workers and all phoretic mites of the colonies sampled in summer 2014 (*N*=58; surviving=24, susceptible=34) were pooled and homogenized for each colony.

#### Individual samples

Adult workers (*N* = 11-13 per colony) and phoretic mites (1-22 per colony) were sampled from 10 surviving and 11 susceptible colonies in autumn 2013. From all three seasons (spring, summer and autumn), honey bee pupae with their corresponding mites (reproductive and non-reproductive) from 47 colonies (8 apiaries), (*N*=29 (3) Surviving and 18 (5) susceptible) were sampled from infested cells and selected for virus analysis.

### Homogenization and RNA extraction

TN buffer (Tris 10mM, NaCl 10 mM; pH 7.6) was added to each sample (25 ml for pooled workers, 100-300 μl for pooled phoretic mites, 250 μl for individual workers and pupae and 100 μl for individual mites) and the sample was homogenized with either a Dispomix® Drive homogenizer (Medic tools) for pooled worker samples or a tissuelyser (Qiagen Retsch MM300, 1 min at 25g/s) for pooled phoretic mites and all individual samples [25]. An aliquot of 50 μl homogenate was used for RNA extraction using the NucleoSpin® RNA II kit, (Macherey-Nagel) following the manufacture’s recommendations. The total extracted RNA was diluted in 60 μl of RNase-free water.

### Reverse transcription, PCR and qPCR assays

The RNA was transcribed to cDNA using M-MLV reverse transcription kit (Promega) following the manufacturer’s recommendations using a defined amount of RNA (1μg for bees and 50 ng for mites, respectively) according to fluorospectrometry (NanoDrop™ 1000) measurements [25]. cDNAs were diluted 10fold in nuclease-free water. With a standard qualitative PCR (0.125 My Taq™ polymerase (Bioline), 5μl 5x buffer, 1 μl of the respective forward and reverse primers (Table S1); 2 min at 95°C, 35 cycles with 20 sec at 95°C, 20 sec at 57°C and 30 sec at 72°C, 2 min at 72°C), pooled worker and phoretic mite samples were screened for the following viruses: Deformed wing virus type A (DWV-A) and type B (DWVB), Acute bee paralysis virus (ABPV), Israeli acute paralysis virus (IAPV), Kashmir bee virus (KBV), Chronic bee paralysis virus (CBPV), Slow bee paralysis virus (SBPV), Sacbrood virus (SBV), Bee Macula-like virus (BeeMLV), Black queen cell virus (BQCV), Lake Sinai virus 1 (LSV1) and Lake Sinai virus 2 (LSV2). Similar, pooled brood samples were screened for DWV-A, DWV-B, ABPV, IAPV, KBV, SBV, LSV1, LSV2, SBPV and CBPV. Positive and negative controls were used for each PCR run. Each PCR Product was analyzed on 1.2 % agarose gel. The agarose gel was stained with GelRed™ and visualized by UV light. With quantitative RT-PCR (RT-qPCR; Kapa SYBR® Fast Master Mix (KAPA, Biosystems), 10 μl master mix, 3 μl cDNA template, 0.4 μl forward and reverse target primers (10 mM) and 6.2 μl Milli-Q water; 3 min at 95°C, 40 cycles of 95°C for 3 sec and 55°C for 30 sec, melting curve: 95°C for 15 sec, 55°C for 15 sec and 95°C for 15 sec) pooled worker samples, where viruses were detected with qualitative PCR, were analyzed to determine virus levels (DWV-A, BQCV, LSV1, LSV2 and SBPV). Individual adult workers and phoretic mites were analyzed individually for DWV-A by use of qPCR (protocol described above). Individual brood samples (pupae and brood mites) were analyzed individually for the viruses detected in the PCR (DWVA, DWV-B, and SBPV, protocol described above). In order to normalize the data according to the amount of RNA, analysis of the *β-actin* gene was performed in parallel for each sample [26]. A Cq cut-off value (according to the value of the negative control) was used to define the disease status (positive or negative).

### Sequencing

To confirm the virus identity of the PCR and qPCR positive samples (pooled and individual), selected PCR-products of each virus were commercially sequenced (Fasteris SA) and compared with reference sequences deposited in GenBank.

### Statistical analyses

Statistical analyses were performed using NCSS Statistical Software [27]. For comparison of virus levels, a two-tailed t-test or a Mann-Whitney U test was done, depending on normal distribution (Kolmogorov-Smirnov, Skewness, Kurtosis and Omnibus Normality). Equally, for comparison of infestation rates among multiple groups one-way ANOVA or a Kruskal Wallis one-way ANOVA on ranks, followed by Dunn-Bonferroni correction was performed. To take account of colony or apiary level variation, the data were additionally run through a linear mixed effects model using R 3.1.2 [28]. Virus values (# copies) were log transformed since the data covered a wide range of values of several magnitudes. Analysis of virus prevalence between groups were done with a Chi-Square test. For all statistical analyses, a significance level of α = 0.05 was applied.

## Results

### Mite population levels

Untreated colonies generally had lower phoretic mite loads (Kruskal-Wallis ANOVA, Dunn’s test, z_spring_>3.04 / z_summer_>2.94, p<0.01, however brood infestation rates were comparable between populations for the sampling period.

### Virus prevalence and viral load

#### Pooled samples

In the pooled worker and phoretic mite samples of the 58 colonies (19 surviving, 39 susceptible) sampled in summer 2014, the following viruses were detected: BQCV, DWV-A, LSV1, LSV2, SBPV and CBPV. All viruses found by PCR were confirmed by sequencing. The surviving colonies had a significantly higher prevalence of BQCV than the susceptible colonies (Chi-Square test: χ^2^=13.44, p<0.01, ~90% of surviving colonies and ~40% of susceptible colonies), but no such differences in prevalence were seen for any of the other tested viruses (Chi-Square tests: χ^2^=1.25-2.80, p>0.05 in all cases, Fig 1).

**Fig 1.**
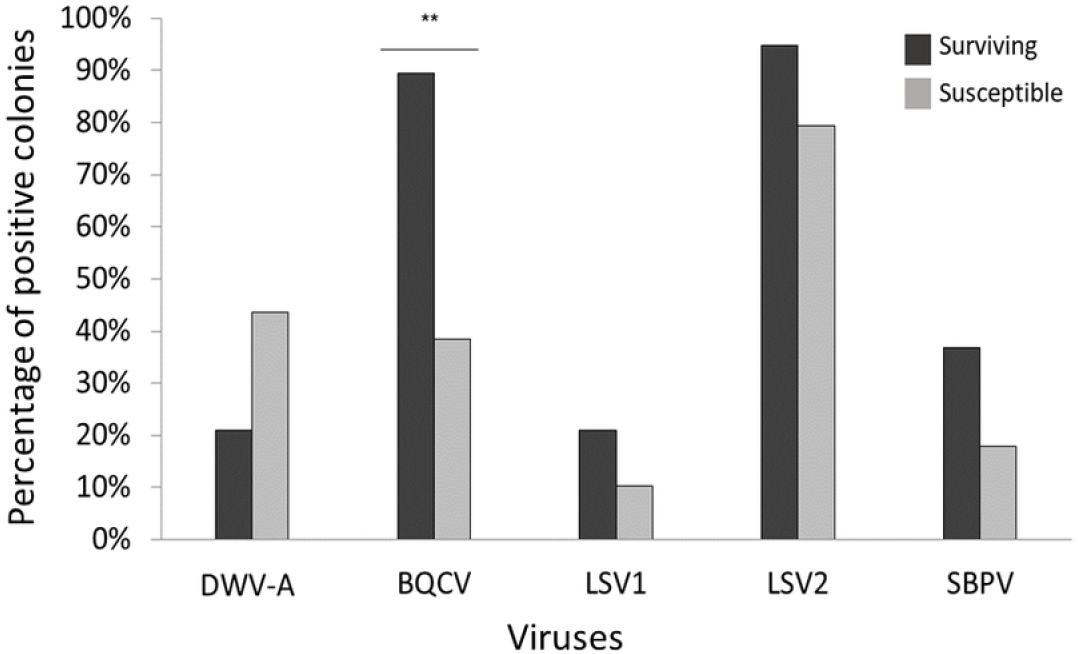
Colony-level Prevalence of the detected viruses in pooled worker samples in summer 2014. While there were no significant differences for most viruses (Chi Square test: χ^2^=1.25-2.81, DWV-A, LSV-1, LSV-2 and SBPV: p>0.05), untreated (surviving) colonies had a significantly higher BQCV prevalence, compared to treated (susceptible) ones (Chi Square test, χ^2^=13.44, ** = p<0.01).

When comparing virus titers between surviving and susceptible colonies, the surviving colonies had higher BQCV and LSV1 loads (BQCV: Mann-Whitney and LSV1: two-tailed t-test; p<0.01), but no significant differences were detected for DWV-A, LSV2 and SBPV (Mann-Whitney, p>0.05 in all cases, Fig 2).

**Fig 2.**
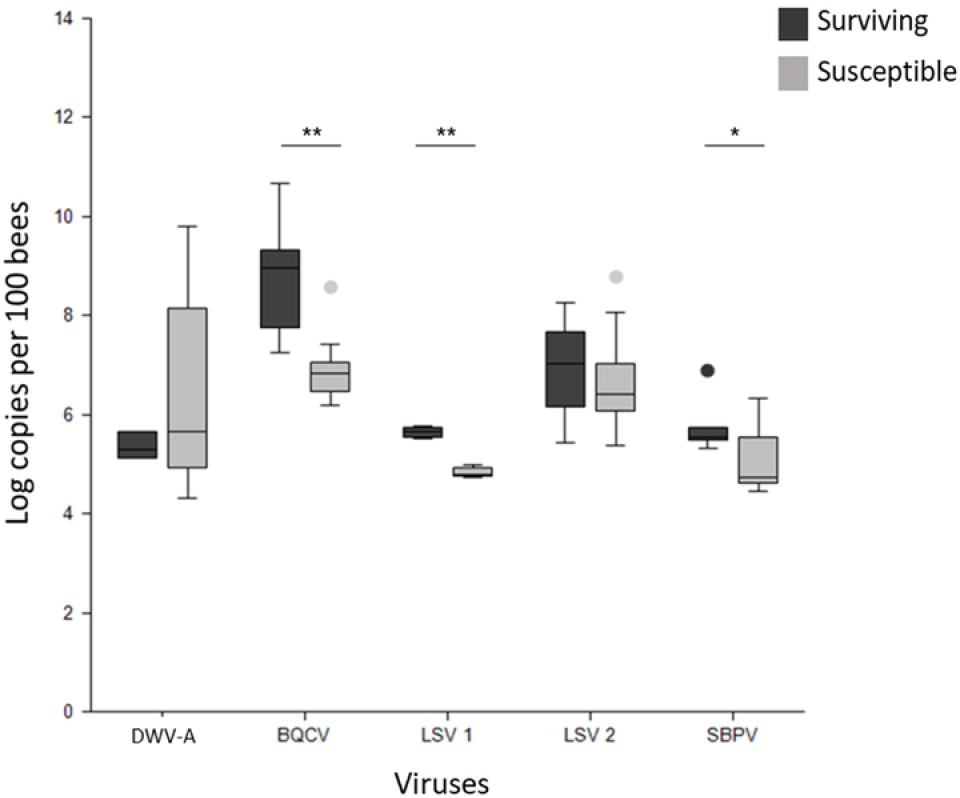
Viral titers of the detected viruses in pooled worker samples of summer 2014 from untreated (surviving) and treated (susceptible) colonies. Medians, interquartile ranges and maxima are shown. While there were no significant differences for most viruses (DWV-A: Mann-Whitney, LSV2, SBPV: two-tailed t-test, p>0.05) surviving colonies had significantly higher BQCV and LSV1 loads, compared to treated ones (BQCV: Mann-Whitney, LSV1: two-tailed t-test; ** = p<0.01).

#### Individual samples

No differences of the DWV-A titers between the surviving and susceptible colonies were found, neither for workers (*N*=201, Mann-Whitney, U_surviving_=4131, U_susceptible_=5463, p>0.05), nor mites (*N*=107, Mann-Whitney, U_surviving_=1395, U_susceptible_=1395, p>0.05). However, surviving colonies showed a significantly lower proportion of DWV-A positive workers than treated colonies did (Chi-Square test: χ^2^=33.751, p<0.01, Fig 3, ~65% surviving workers and ~95% of susceptible workers).

**Fig 3.**
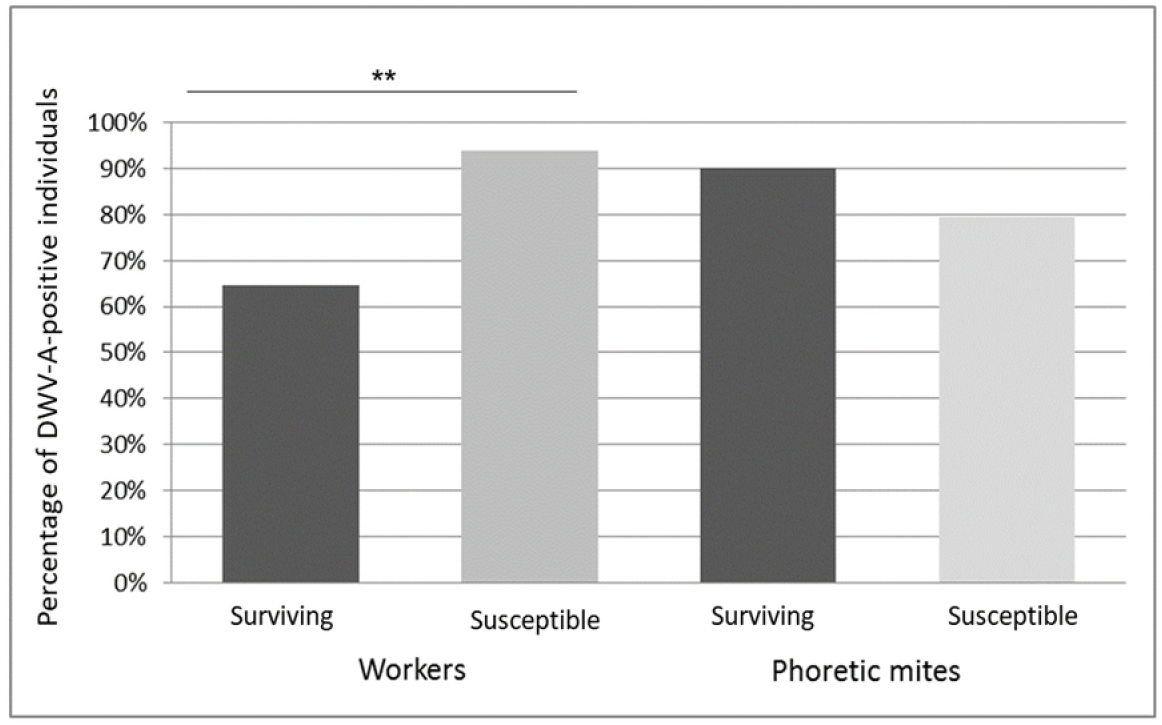
Prevalence of DWV-A in workers and phoretic mites of autumn 2013 from untreated (surviving) and treated (susceptible) colonies. Means are shown. While there was no significant difference in the proportions of DWV-A positive mites (Chi Square, χ^2^=2.455, p>0.05), significantly fewer workers from surviving colonies had detectable DWV-A titers compared to susceptible colonies (Chi Square, χ^2^ 33.751, ** = p<0.01).

Viruses detected in brood and associated mites were Deformed wing virus-A (DWV-A), Deformed wing virus-B (DWV-B) and Slow bee paralysis virus (SBPV). No significant differences of virus prevalence were detected between susceptible and surviving pupae and mites for all tested viruses (Chi-Square: χ^2^=0.76-3.17, p>0.05). Comparison of titers from susceptible and surviving pupae revealed that those from the susceptible colonies had significantly higher DWV-A titers (Mann-Whitney, workers: U_treated_ = 1511, U_surviving_ = 541, mites: U_susceptible_ = 2500, U_surviving_ = 620, p<0.01 for workers and mites, Fig 4).

**Fig 4.**
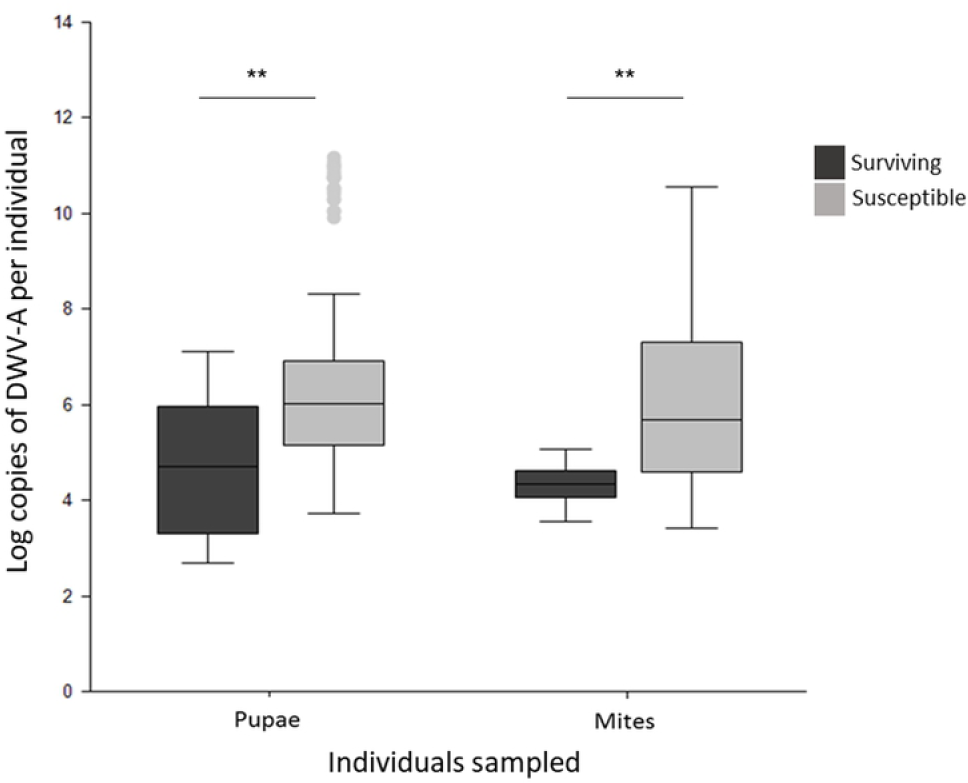
DWV-A titers of individual honeybee pupae and their associated mites in untreated (surviving) and treated (susceptible) colonies from all seasons. Medians, interquartile ranges and maxima are shown. Both pupae and mites from susceptible colonies had significantly higher DWV-A titers than those from surviving colonies (Mann-Whitney, workers: U_susceptible_=1511, U_surviving_=541, mites: U_susceptible_=2500, U_surviving_=620, ** = p<0.01).

No significant differences were detected between surviving and susceptible pupae and mites for DWV-B and SBPV titers.

## Discussion

The proportion of adult workers and pupae that tested positive for one of the most prominent mite-transmitted viruses (DWV-A) was lower in surviving colonies than in susceptible controls, meaning the virus may have had a reduced transmission frequency, likely brought about by fewer phoretic mites. Individual adult workers in the surviving population had a higher prevalence for BQCV and higher titers of this virus as well as one other not commonly associated with *V. destructor* (LSV1) and this may suggest a reduced ability to overcome infections from these viruses.

Though the DWV-A titer level was not different between individual adult worker bees in both colony groups (surviving and susceptible) the proportion of workers that tested positive for DWV-A was significantly lower. Phoretic mites did not have significantly different viral prevalence or titer between surviving and susceptible groups, meaning the mites were just as capable of spreading infection in surviving colonies as susceptible colonies. If viral tolerance played a role in mite-survivability we may not have seen such a distinction in viral prevalence. As prevalence is reduced in surviving colonies, it can be considered that the control factor is focused on reducing the probability of viral infection, i.e. reducing the mite loads. Viral prevalence in infested pupae was not different between surviving and susceptible colony groups and this also aligns with the idea of mite-targeted survival strategies: If a cell is infested, it has the same probability of becoming infected with a mite-transmitted virus.

The reason we do not see this lower prevalence in the pupae could be because only infested pupae were sampled, though no differences in pupal infestation rates were found in this study, there were differences in phoretic mite load, and these surviving colonies have historically displayed consistently lower mite loads both phoretically and in brood [8].

Interestingly, there was a difference in DWV-A titers in both pupae and associated mites when comparing the colony groups (surviving and susceptible): surviving pupae and mites had lower titers of this virus. There is a possibility that this is due to the lower frequency of bees being bitten by mites (due to reduced mite load) and therefore a lower general level of DWV-A circulating within the population, however workers that did test positive for the virus did not display a difference in titer, nor did the phoretic mites. More likely, the mite-surviving strategy relies on reducing the number of offspring produced in mite-infested pupal cells (SMR: [7,8,29] and the lower number of offspring reduces viral transference in the closed system of the cell. These differences, though statistically significant, were small, but they provide further evidence that reducing mite loads is the predominant surviving trait.

Surviving colonies had higher titers of BQCV and LSV1 and a higher prevalence of BQCV. In another surviving population in Sweden, BQCV and SBV titers decreased substantially compared to a local susceptible population [21]. The history of these Norwegian colonies contains a sharp reduction in population and a steady increase again from those few colonies that survived [7]. It is possible that the reduction of genetic material being bred from at the time created a larger susceptibility to generally non-lethal threats as has been shown in previous work [30]. Despite evolved strategies to combat inbreeding [31] bee species can suffer the effects of genetic bottlenecking [32–35]. It may also be possible that the strategy of mite-survival may leave the colonies more vulnerable to other pathogens, in the way it is employed.

The presence of SMR in all recorded surviving populations [7,8,29] is good evidence to suggest that the reduction of mite infestation levels is the most successful natural strategy to mitigate the damages of *V. destructor* in populations of Western honey bee. With consistent evidence of hindering mite population growth in all populations, this strategy seems to be present everywhere Western honeybees are permitted to adapt naturally and should therefore be a core focus of *Varroa destructor*-resistant breeding efforts. Viral tolerance cannot be discounted, however future studies on mite survivability might benefit from a focus on regulating the parasite populations and not enduring them.

## Acknowledgments

We are grateful to the cooperating beekeepers that allowed us to collect samples from their honeybee colonies.

